## Supplemental Figures for "Neurovascular coupling and bilateral connectivity during NREM and REM sleep"

#### Supplemental video captions

**Video S1-3 | Measurements of arousal and associated changes in neural activity and hemodynamics.** Whisker motion, IOS reflectance, and eye camera activity are shown alongside measurements of neural activity and hemodynamics. **Video S1**, The awake state shows a large amount of whisker motion and elevations in heart rate. The eye is open. High frequency neural activity increases and low frequency activity decreases during whisking events, with corresponding decreases in reflectance. An increase in blood volume/[HbT] corresponds to a decrease in pixel reflectance, as more light is absorbed by the increase in hemoglobin. **Video S2**, The NREM state shows little whisker motion and a lower heart rate. The eye is still open. Low frequency (delta band) cortical neural activity is elevated, and there are large changes in reflectance. **Video S3**, The transition into the REM state shows an increase in whisker motion and an increase heart rate, similar to the awake state. The eye remains open. The neural activity in the hippocampal theta band and upper frequencies of cortical neural activity increase while [HbT] increases substantially.

#### Supplemental figure captions

**Fig. S1 | Localization of electrodes and hemodynamic regions of interest. a**, Image of a cortical window showing cortical vasculature. **b**, Peak pixel-wise cross-correlation (1-2 second lag) between gamma band power [30-100 Hz] and pixel reflectance during the first 60 minutes of data. **c**, same as **b** for multi-unit activity (300-3000 Hz). The peak cross-correlation was used to localize a 1mm diameter region of interest (ROI). **d**, Histological example of a coronal section stained with cytochrome oxidase (CO). Small holes in the slice indicate the location of cortical stereotrodes in the vibrissa barrel cortex. **e,** Histological example of a coronal section stained with CO. Electrode path is visible and terminates in the CA1 region of hippocampus.

**Fig. S2 | Whisker stimulation causes increases in neural activity and blood volume.** Comparisons between contralateral, ipsilateral, and auditory whisker stimulation and the corresponding changes in vibrissa cortical MUA/LFP, hippocampal MUA/LFP, and hemodynamic changes (reflectance/[HbT]) (n = 14 mice, 28 hemispheres). (**a-c**) Contralateral whisker stimulation caused large increases in vibrissa cortical MUA power (78.1 ± 66.7%) in comparison to ipsilateral (30.8 ± 37.5%) and auditory (13.7 ± 16.8%) stimulation. (**d-f**) LFP gamma band power increased (110.4 ± 96.7%) in comparison to ipsilateral (36.8 ± 32.8%) and auditory stimulation (17.4 ± 15.7%). (**g-i**) All three forms of stimulation caused relatively similar changes in hippocampal MUA power (contralateral: 60.1 ± 34.3%, ipsilateral: 57 ± 33.4%, auditory: 42 ± 41.8%) and in (**j-r**) LFP gamma band power (contralateral: 32.5 ± 17.2%, ipsilateral: 28 ± 14.3%, auditory: 18.6 ± 8.1%). Contralateral whisker stimulation caused increases in total hemoglobin Δ[HbT] (16.8 ± 5.1 μM corresponding to a -2.4 ± 0.7% reflectance) larger than those from ipsilateral (9 ± 3.5 μM, -1.3 ± 0.5%) and auditory (4.8 ± 2.3 μM, -0.7 ± 0.3%) stimulation.

**Fig. S3 | Volitional whisking causes increases in neural activity and hemodynamics.** Changes in vibrissa cortical MUA/LFP, hippocampal MUA/LFP, and blood volume during whisking events of various durations (0.5-2 seconds, 2-5 seconds, > 5 seconds) (n = 14 mice, 28 hemispheres). (**a-c**) Extended whisking caused larger increases in vibrissa cortical MUA power (16.5 ± 10.8%) in comparison to moderate (9.7 ± 7%) and brief (5 ± 4.8%) durations. (**d-f**) LFP gamma band power was higher during extended whisking (19.2 ± 18.9%) in comparison to moderate (15.1 ± 16.3%) and brief (6.7 ± 9.2%) durations. (**g-i**) Whisking drives increases in hippocampal MUA power (brief: 5.1 ± 5%, moderate: 12 ± 7.7%, extended: 16.9 ± 10.1%) and in (**j-r**) LFP gamma band power (brief: 5.7 ± 11.7%, moderate: 13.9 ± 14.3%, extended: 18.4 ± 14.9%). Extended whisking caused increases in total hemoglobin Δ[HbT] (12.1 ± 5.6 μM corresponding to a -1.7 ± 0.8% reflectance) that were larger than those seen in moderate (7.1 ± 3.3 μM, -1 ± 0.5%) and brief (2.5 ± 1.4 μM, -0.4 ± 0.2%) duration whisking events.

**Fig. S4 | Amplitude of hemodynamic oscillations are largest during NREM and REM sleep.** **a**, The peak-to-peak amplitude of Δ[HbT] oscillations during awake rest (32.3 ± 4.4 μM) were significantly smaller than those during contiguous NREM (87.3 ± 9.9 μM, GLME, $p < 9\times10^{-32}$) and contiguous REM (142.1 ± 20.7 μM, GLME, $p < 1.5\times10^{-53}$) sleep. **b**, Mean peak Δ[HbT] of individual awake resting events (17 ± 3.2 μM) were significantly smaller than the peaks during contiguous NREM (69.8 ± 10.7 μM, GLME, $p < 1.1\times10^{-37}$) and contiguous REM (107.7 ± 13.3 μM, GLME, $p < 3.5\times10^{-55}$) sleep (n = 14 mice, 28 hemispheres). **c**, Peak-to-peak amplitude of ΔD/D oscillations during awake rest (16.6 ± 4 μM) were significantly smaller by those during contiguous NREM (38 ± 15.8%, GLME, $p < 3.6\times10^{-10}$) and contiguous REM (59.9 ± 15 μM, GLME, $p < 3.3\times10^{-16}$) sleep. **d**, Mean peak ΔD/D of individual awake resting events (7.4 ± 2.6 μM) were significantly smaller than the peaks during contiguous NREM (26.4 ± 10.1 μM, GLME, $p < 1.9\times10^{-14}$) and contiguous REM (49.9 ± 9.1 μM, GLME, $p < 2\times10^{-24}$) sleep (awake rest: n = 6 mice, 29 arterioles**,** contiguous NREM: n = 6 mice, 21 arterioles, contiguous REM: n = 5 mice, 10 arterioles). *p < 0.05, **p < 0.01, ***p < 0.001 GLME.

**Fig. S5-8 | Sleep drives hemodynamic fluctuations larger than awake behaviors.** Examples showing the hemodynamic and neural changes accompanying transitions among the NREM, REM and awake states. **a**, Plot of nuchal muscle EMG power and body motion via a pressure sensor located beneath the mouse. **b**, Plot of the whisker position and heart rate. **c**, Changes in total hemoglobin Δ[HbT] within the ROIs over the putative vibrissa cortex. Inset shows images of the two windows and respective ROIs. **d**, Normalized left vibrissa cortex LFP power (ΔP/P). **e**, Normalized right vibrissae cortex LFP power. **f**, Normalized CA1 LFP power.

**Fig. S9 | Behavioral measurements demarcate transitions between arousal states.** **a**, Power in rfc-Awake electromyograph (EMG) recordings (1.3 ± 1.3 from the nuchal muscles were substantially smaller during rfc-NREM sleep (0.25 ± 2, GLME, $p < 3.2\times10^{-12}$) and during rfc-REM sleep (0.05 ± 1.6, GLME, $p < 1.7\times10^{-21}$) in comparison to the rfc-Awake state. **b**, Variance in the whisker angle during rfc-NREM (2.4 ± 1.5 $deg^2$, GLME, $p < 1.3\times10^{-12}$) was significantly less than that of the awake state (25.7 ± 9 $deg^2$). Whisker angle variance during rfc-REM (18.6 ± 7.3 $deg^2$, GLME, p < 0.003), though statistically different, was much more similar to the awake state due to mice sporadically moving their whiskers during rfc-REM sleep, analogous to rapid-eye movement seen in humans. **c**, Heart rate during the rfc-Awake state was 7.5 ± 0.7 Hz. During rfc-NREM sleep, the heart rate dropped to 6.1 ± 0.6 Hz (GLME, $p < 1.8\times10^{-13}$), and was elevated slightly during rfc-REM to 7.1 ± 0.5 Hz (GLME, p < 0.004) (n = 14 mice). *p < 0.05, **p < 0.01, ***p < 0.001 GLME.

**Fig. S10 | Volitional whisking causes arteriole dilation.** **a**, Brief awake whisking events (0.5-2 seconds) spurred a small dilation 0.8 ± 0.9%. **b**, Moderate length awake whisking events (2-5 seconds) lead to a more robust dilation (8.3 ± 4.9%). **c**, Extended awake whisking events (> 5 seconds) produced the largest dilations (10.9 ± 4.3%, n = 29 arterioles).

**Fig. S11-14 | Arteriole dilatations during sleep are larger than those during the awake state.** Example showing the vascular and neural changes accompanying transitions among the NREM, REM and awake states. Arterial diameters were imaged using two-photon microscopy. **a**, Nuchal muscle activity through normalized EMG and body motion via a pressure sensor located beneath the mouse. **b**, Whisker position. **c**, Changes in arteriole diameter $\Delta D/D$ (%) in the putative vibrissa cortex. **d**, Normalized LFP power from the left hemisphere stereotrode located in the left vibrissa cortex. **e**, Normalized LFP power from the stereotrode in left CA1.

**Fig. S15 | Transitional changes in hemodynamics are consistent across each day.** Average change in total hemoglobin $\Delta$[HbT] for different days for various behavioral state transitions. Each colored line indicates a unique day of imaging (n = 14 mice). **a**, Transition from rfc-Awake to rfc-NREM. **b**, Transition from rfc-NREM to rfc-Awake. **c**, Transition from rfc-NREM to rfc-REM. **d**, Transitions from rfc-REM to rfc-Awake.

**Fig. S16 | Isoflurane drives larger vasodilations than sleep.** Example showing the blood volume and neural changes accompanying transitions between NREM and awake states followed by administration of isoflurane. Isoflurane is a potent vasodilator and causes the cortical vasculature to reach a maximum or near-maximum saturation in blood volume. **a**, Plot of nuchal muscle EMG power and body motion via a pressure sensor located beneath the mouse. **b**, Plot of the whisker position and heart rate. **c**, Changes in total hemoglobin $\Delta$[HbT] within the ROIs over the putative vibrissa cortex. Inset shows images of the two windows and respective ROIs. **d**, Normalized left vibrissae cortex LFP power ($\Delta P/P$). **e**, Normalized right vibrissae cortex LFP power. **f**, Normalized CA1 LFP power. **g**, Average change in total hemoglobin $\Delta$[HbT] within the ROI during different arousal states. Circles represent individual hemispheres of each mouse and diamonds represent population averages, with error bar showing ± 1 standard deviation (n = 14 mice, 28 hemispheres; Isoflurane: n = 9, 18 hemispheres). $*p < 0.05$, $**p < 0.01$, $***p < 0.001$ GLME.

**Fig. S17 | Arousal state dependence of low-frequency neural power and coherence[2].** Spectral power and MOC2 at 0.1 or 0.01 Hz during different arousal states. **a-h** n = 14 mice (n*2 hemispheres in **a,b,e,f**) for all arousal states except Alert: n = 12 mice, Asleep: n = 13 mice. **a-d**, Gamma band power. **e-h**, $\Delta$[HbT]. **i,j** Spectral power at 0.1 or 0.01 Hz during different arousal states for arteriole $\Delta D/D$ (Rest: n = 6 mice, 29 arterioles, NREM: n = 6 mice, 21 arterioles, REM: n = 5 mice, 10 arterioles, Awake: n = 6 mice, 27 arterioles, All data: n = 6 mice, 29 arterioles). Circles represent individual hemispheres of each mouse (**a-h**) or individual arterioles (**i,j**) and diamonds represent population averages, with error bar showing ± 1 standard deviation. Data is presented in **Table 1-5**. $*p < 0.05$, $**p < 0.01$, $***p < 0.001$ GLME.

**Fig. S18 | Correlations in neural activity between hemispheres increase during sleep. a**, Mean delta band power spectral density during different arousal states. **b**, Mean coherence (between hemispheres) in the changes in the envelope (≤ 1 Hz) of delta band power [1-4 Hz] between left and right vibrissa cortex during different arousal states. **c**, Average delta band power Pearson's correlation coefficient between left and right vibrissa cortex during different arousal states. Circles represent individual mice and diamonds represent population averages ± 1 standard deviation. **d-f,** Same as in a-c except for the theta band power [4-10 Hz]. **g-i,** Same as in **a-c** except for the alpha band power [10-13 Hz]. **j-l,** Same as in **a-c** except for the beta band power [13-30 Hz]. MoC2 between the left and right somatosensory cortex for each LFP band during each arousal state exceeded the 95% confidence level for all envelope frequencies below 1 Hz. **a-l** n = 14 mice (n*2 hemispheres in **a,d,g,j**) for all arousal states except Alert: n = 12 mice, Asleep: n = 13 mice. Data is presented in **Table S3-S7**. *p < 0.05, **p < 0.01, ***p < 0.001 GLME.

**Fig. S19 | Arousal state dependence of low-frequency neural power and coherence.** Spectral power and MOC2 at 0.1 or 0.01 Hz for different arousal states. **a-p** n = 14 mice (n*2 hemispheres in **a,b,e,f,i,j,m,n**) for all arousal states except Alert: n = 12 mice, Asleep: n = 13 mice. **a-d**, Delta band power. **e-h**, Theta band power. **i-l**, Alpha band power. **m-p**, Beta band power. Circles represent individual hemispheres of each mouse and diamonds represent population averages, with error bar showing ± 1 standard deviation. Data is presented in **Table S3-S7**. *p < 0.05, **p < 0.01, ***p < 0.001 GLME.

**Fig. S20 | Arousal state dependence of low-frequency neural-hemo coherence (for Fig. 8).** Coherence between LFP and changes total hemoglobin Δ[HbT] evaluated at 0.1 and 0.01 Hz. **a**, Mean coherence between the envelope (≤ 1 Hz) of delta band power [1-4 Hz] and Δ[HbT] in a single cortical hemisphere during different arousal states at 0.1 (**b**) and 0.01 (**c**) Hz. **d-f**, Same as in **a-c** except for theta band power [4-10 Hz]. **g-i**, Same as in **a-c** except for alpha band power [10-13 Hz]. **j-l**, Same as in **a-c** except for beta band power [13-30 Hz]. **m-o**, Same as in **a-c** except for gamma band power [30-100 Hz]. **a-o** n = 14 mice (n*2 hemispheres) for all arousal states except Alert: n = 12 mice, Asleep: n = 13 mice. Data is presented in **Table S8**. *p < 0.05, **p < 0.01, ***p < 0.001 GLME.

**Fig. S21 | Correction of slow drifts in reflectance during IOS imaging. a**, Image of bilateral hemispheres during IOS imaging with localized left, right ROIs as well as a region over the central cement that is used to correct a slow exponential drift in the camera's sensitivity. The drift in reflectance of the cement (orange) is fit with an exponential (purple) and is used to correct the drifts in the lateral ROIs. **b**, Raw pixel reflectance from the left hemisphere ROI. The exponential drift is clearly visible prior to correction. **c**, Raw pixel reflectance from the right hemisphere ROI. **d**, The exponential drift from the cement ROI is inverted and normalized to correct pixel reflectance in each hemisphere. **e**, Original (red) vs. corrected (purple) pixel reflectance for the left hemisphere ROI. **f**, Original (blue) vs. corrected (purple) pixel reflectance for the right hemisphere ROI.

**Fig. S22 | Random forest model validation.** All data from the first and last day of imaging from each animal was manually scored as rfc-Awake, rfc-NREM, or rfc-REM. Alternating 15-minute periods of data from these two days were divided into two discrete sets: one for model training, the other is held back for model validation beyond the out-of-bag error obtained from the training data set. A confusion matrix containing each IOS animal's (n = 14) random forest model predictions of the held back, second data set compared to its manual scores are presented in **a**. The total model accuracy across all 14 animals was 91.3%, with the most accurate predictions coming from the most prevalent classification class (rfc-Awake), followed by rfc-NREM and then rfc-REM.

**Table S1 | Duration of each arousal state from each animal used in IOS experiments.**

| **Animal ID** | **Total Data (Hours)** | **Awake Data (Hours)** | **Awake Data (% of Total)** | **NREM Data (Hours)** | **NREM Data (% of Total)** | **REM Data (Hours)** | **REM Data (% of Total)** |
|---|---|---|---|---|---|---|---|
| ***T99*** | 19.8 | 6.4 | 32.2 | 11.4 | 57.7 | 2.0 | 10.1 |
| ***T101*** | 18.8 | 12.6 | 66.8 | 5.2 | 27.6 | 1.1 | 5.6 |
| ***T102*** | 20.3 | 12.9 | 63.4 | 6.3 | 31.0 | 1.1 | 5.6 |
| ***T103*** | 21.0 | 15.2 | 72.4 | 4.8 | 22.9 | 1.0 | 4.7 |
| ***T105*** | 20.5 | 12.4 | 60.5 | 7.8 | 38.0 | 0.3 | 1.4 |
| ***T108*** | 18.8 | 12.2 | 65.0 | 6.0 | 32.1 | 0.5 | 2.9 |
| ***T109*** | 19.8 | 16.0 | 81.0 | 3.4 | 17.1 | 0.4 | 1.9 |
| ***T110*** | 16.0 | 13.0 | 81.5 | 2.6 | 16.4 | 0.3 | 2.1 |
| ***T111*** | 21.3 | 12.2 | 57.5 | 7.5 | 35.3 | 1.5 | 7.2 |
| ***T119*** | 29.5 | 19.4 | 65.8 | 7.1 | 24.1 | 3.0 | 10.1 |
| ***T120*** | 28.8 | 13.8 | 47.9 | 13.6 | 47.1 | 1.4 | 5.0 |
| ***T121*** | 27.5 | 10.2 | 37.2 | 15.2 | 55.4 | 2.0 | 7.4 |
| ***T122*** | 27.3 | 7.0 | 25.7 | 18.4 | 67.5 | 1.9 | 6.8 |
| ***T123*** | 32.8 | 10.9 | 33.2 | 21.5 | 65.5 | 0.4 | 1.3 |

**Table S2 | Duration of each arousal state from each animal used in two photon experiments.**

| **Animal ID + Arteriole ID** | **Total Data (Minutes)** | **Baseline Diameter (μm)** | **Awake Data (Minutes)** | **NREM Data (Minutes)** | **REM Data (Minutes)** |
|---|---|---|---|---|---|
| ***T115 A1*** | 105.0 | 30.6 | 88.9 | 13.3 | 2.8 |
| ***T115 A2*** | 120.0 | 24.5 | 105.7 | 14.3 | 0.0 |
| ***T115 A3*** | 120.1 | 12.7 | 109.3 | 10.8 | 0.0 |
| ***T115 A4*** | 30.0 | 23.5 | 28.4 | 1.6 | 0.0 |
| ***T115 P1*** | 90.0 | 16.3 | 88.2 | 1.8 | 0.0 |
| ***T116 A1*** | 90.1 | 24.9 | 55.3 | 34.8 | 0.0 |
| ***T116 A2*** | 150.0 | 22.8 | 106.7 | 43.3 | 0.0 |
| ***T116 A3*** | 105.0 | 20.8 | 88.8 | 16.2 | 0.0 |
| ***T117 A1*** | 90.1 | 26.3 | 70.8 | 18.1 | 1.2 |
| ***T117 A2*** | 60.0 | 18.5 | 52.3 | 7.7 | 0.0 |
| ***T117 A3*** | 75.0 | 15.9 | 55.2 | 19.8 | 0.0 |
| ***T117 A4*** | 15.0 | 17.7 | 14.2 | 0.8 | 0.0 |
| ***T118 A1*** | 30.0 | 22.6 | 13.6 | 13.8 | 2.6 |
| ***T118 A2*** | 60.0 | 19.0 | 42.2 | 13.8 | 4.0 |
| ***T118 A3*** | 90.0 | 20.6 | 58.1 | 27.6 | 4.3 |
| ***T118 P1*** | 15.0 | 27.3 | 13.4 | 1.6 | 0.0 |

| | | | | | |
|---|---|---|---|---|---|
| ***T125 A1*** | 60.0 | 22.5 | 54.1 | 5.9 | 0.0 |
| ***T125 A2*** | 75.1 | 19.3 | 60.8 | 14.3 | 0.0 |
| ***T125 A3*** | 45.0 | 17.5 | 43.2 | 1.8 | 0.0 |
| ***T125 A4*** | 75.0 | 17.7 | 58.0 | 15.0 | 2.0 |
| ***T125 A5*** | 75.0 | 17.9 | 63.3 | 10.1 | 1.6 |
| ***T125 P1*** | 30.1 | 14.6 | 18.2 | 9.3 | 2.6 |
| ***T126 A1*** | 75.1 | 31.5 | 72.8 | 2.3 | 0.0 |
| ***T126 A2*** | 75.0 | 23.7 | 73.6 | 1.4 | 0.0 |
| ***T126 A3*** | 90.0 | 25.2 | 88.2 | 1.8 | 0.0 |
| ***T126 A4*** | 105.0 | 21.8 | 93.1 | 10.6 | 1.3 |
| ***T126 A5*** | 60.0 | 18.6 | 46.3 | 12.1 | 1.6 |
| ***T126 A6*** | 45.0 | 18.3 | 44.3 | 0.7 | 0.0 |
| ***T126 P1*** | 45.0 | 12.0 | 42.4 | 2.6 | 0.0 |

**Table S3 | IOS arousal state classification criteria**

| **Arousal state** | **Color** | **Duration** | **Origin** | **Criterion** |
|---|---|---|---|---|
| rfc-Awake | Light black | 5 seconds | Random forest classifier | rfc-Awake periods were denoted by moderate cortical gamma band power, low cortical delta band power, moderate-high whisker motion, moderate-high heart rate, and high EMG power. |
| rfc-NREM | Cyan | 5 seconds | Random forest classifier | rfc-NREM periods were denoted by moderate cortical gamma band power, high cortical delta band power, little-no whisker motion, low heart rate, and low EMG power. |
| rfc-REM | Dark red | 5 seconds | Random forest classifier | rfc-REM periods were denoted by high cortical gamma band power, high hippocampal theta band power, moderate whisker motion, moderate heart rate, and very low EMG power. |
| Awake Rest | Green | ≥ 10 seconds | Subsets of rfc-Awake | Large epochs (typically > 60 seconds) were manually verified to be truly awake. Awake Rest was defined as no body movement or whisker motion and occurred at least 5 seconds away from a whisker stimulation. |
| Awake Whisking | Dark blue | 0-5 seconds post whisk | Subsets of rfc-Awake | Large epochs (typically > 60 seconds) were manually verified to be truly awake. Awake Whisking was defined as whisking motion that lasted between 2 and 5 seconds long and occurred at least 5 seconds away from a whisker stimulation. |
| Awake Stimulation | Pink | 1-2 seconds post stim | Subsets of rfc-Awake | Large epochs (typically > 60 seconds) were manually verified to be true awake behavior. Awake Stimulation was defined as a directed air puff (0.1 seconds, 10 PSI) to a contralateral whisker pad. |
| Contiguous NREM | Purple | ≥ 30 seconds | Subsets of rfc-NREM' | rfc-NREM periods that meet duration threshold and occurred in the absence of whisker stimulation |
| Contiguous REM | Orange | ≥ 60 seconds | Subsets of rfc-REM | rfc-REM periods that meet duration threshold and occurred in the absence of whisker stimulation. Long REM events that were broken up by ≤ 10 seconds of misclassification were linked as a single event. |
| Alert | Gold | 15 minutes | Random forest classifier | Full 15-minute recordings with ≥ 80% rfc-Awake classifications and lacked whisker stimulation. |
| Asleep | Light blue | 15 minutes | Random forest classifier | Full 15-minute recordings with ≥ 80% rfc-NREM or rfc-REM classifications and lacked whisker stimulation. i.e. ≤ 20% rfc-Awake classification. |

| | | | | |
|---|---|---|---|---|
| All Data | Brown | 15 minutes | Random forest classifier | All 15-minute recordings that lacked whisker stimulation regardless of rfc classifications. |

**Table S4 | 2-photon arousal state classification criteria**

| Arousal state | Color | Duration | Origin | Criterion |
|---|---|---|---|---|
| m-Awake | Light black | 5 seconds | Manually classified | m-Awake periods were denoted by moderate cortical gamma band power, low cortical delta band power, moderate-high whisker motion, moderate-high heart rate, and high EMG power. |
| m-NREM | Cyan | 5 seconds | Manually classified | m-NREM periods were denoted by moderate cortical gamma band power, high cortical delta band power, little-no whisker motion, low heart rate, and low EMG power. |
| m-REM | Dark red | 5 seconds | Manually classified | m-REM periods were denoted by high cortical gamma band power, high hippocampal theta band power, moderate whisker motion, moderate heart rate, and very low EMG power. |
| Awake Rest | Green | ≥ 10 seconds | Subsets of m-Awake | Epochs were manually verified to be truly awake. Awake Rest was defined as no body movement or whisker motion. |
| Awake Whisking | Dark blue | 0-5 seconds post whisk | Subsets of m-Awake | Epochs were manually verified to be truly awake. Awake Whisking was defined as whisking motion that lasted between 2 and 5 seconds long. |
| Contiguous NREM | Purple | ≥ 30 seconds | Subsets of m-NREM' | m-NREM periods that meet duration threshold. |
| Contiguous REM | Orange | ≥ 60 seconds | Subsets of m-REM | m-REM periods that meet duration threshold. |
| Alert | Gold | 15 minutes | Manually classified | Full 15-minute recordings with ≥ 80% m-Awake classifications. |
| All Data | Brown | 15 minutes | Manually classified | All 15-minute recordings regardless of classification (entire data set). |

**Table S5 | Spectral power in delta band, theta band, alpha band, and beta band at 0.1 Hz**

| Spectral Power at 0.1 Hz | Awake Rest | Cont. NREM | Cont. REM | Alert | Asleep | All Data |
|---|---|---|---|---|---|---|
| Delta band Power (a.u.) | 0.9 ± 0.1 | 4.7 ± 3.2 ($p < 4.1\times10^{-10}$) | 3.5 ± 1.8 ($p < 1.2\times10^{-5}$) | 1.1 ± 0.4 ($p < 0.8$) | 5 ± 3.3 ($p < 2.9\times10^{-11}$) | 3 ± 1.7 ($p < 0.0003$) |
| Theta band Power (a.u.) | 0.9 ± 0.1 | 19.2 ± 59.9 ($p < 0.08$) | 27.6 ± 72.6 ($p < 0.01$) | 1 ± 0.3 ($p < 0.99$) | 2.5 ± 1.7 ($p < 0.88$) | 1.7 ± 0.6 ($p < 0.94$) |
| Alpha band Power (a.u.) | 0.9 ± 0.1 | 59.8 ± 177 ($p < 0.008$) | 47.3 ± 100 ($p < 0.04$) | 1.2 ± 0.4 ($p < 0.99$) | 6.5 ± 4.2 ($p < 0.8$) | 3.9 ± 1.9 ($p < 0.89$) |
| Beta band Power (a.u.) | 0.9 ± 0.1 | 123.9 ± 397.6 ($p < 0.01$) | 69.6 ± 170.3 ($p < 0.15$) | 1.3 ± 0.6 ($p < 0.99$) | 7.7 ± 4.8 ($p < 0.89$) | 4.6 ± 2.5 ($p < 0.94$) |

*Mean ± 1 standard deviation. p-values as a comparison to "Rest".*

**Table S6 | Spectral power in delta band, theta band, alpha band, and beta band at 0.01 Hz**

| **Spectral Power at 0.01 Hz** | **Alert** | **Asleep** | **All Data** |
|---|---|---|---|
| Delta band Power (a.u.) | 8.3 ± 6 | 21.3 ± 21.3 ($p < 0.004$) | 18.7 ± 16.3 ($p < 0.02$) |
| Theta band Power (a.u.) | 5.5 ± 2.7 | 20.9 ± 18.3 ($p < 8.6\times10^{-6}$) | 13.8 ± 7.5 ($p < 0.01$) |
| Alpha band Power (a.u.) | 6.7 ± 4.5 | 62.5 ± 40.7 ($p < 3.9\times10^{-10}$) | 41.2 ± 24.8 ($p < 2.2\times10^{-5}$) |
| Beta band Power (a.u.) | 10.8 ± 10 | 91.4 ± 60.9 ($p < 6.9\times10^{-10}$) | 62.7 ± 35.9 ($p < 1.5\times10^{-5}$) |

*Mean ± 1 standard deviation. p-values as a comparison to "Alert".*

**Table S7 | Magnitude of $Coherence^2$ of bilateral delta band, theta band, alpha band, and beta band at 0.1 Hz**

| **$Coherence^2$ at 0.1 Hz** | **Awake Rest** | **Cont. NREM** | **Cont. REM** | **Alert** | **Asleep** | **All Data** |
|---|---|---|---|---|---|---|
| Delta band ($Coherence^2$) | 0.06 ± 0.06 | 0.07 ± 0.06 ($p < 0.59$) | 0.03 ± 0.03 ($p < 0.03$) | 0.11 ± 0.08 ($p < 0.002$) | 0.08 ± 0.06 ($p < 0.37$) | 0.07 ± 0.05 ($p < 0.63$) |
| Theta band ($Coherence^2$) | 0.05 ± 0.04 | 0.08 ± 0.07 ($p < 0.08$) | 0.09 ± 0.06 ($p < 0.03$) | 0.14 ± 0.09 ($p < 1.2\times10^{-5}$) | 0.07 ± 0.06 ($p < 0.18$) | 0.08 ± 0.05 ($p < 0.13$) |
| Alpha band ($Coherence^2$) | 0.02 ± 0.02 | 0.19 ± 0.06 ($p < 8.7\times10^{-17}$) | 0.1 ± 0.05 ($p < 6.1\times10^{-6}$) | 0.09 ± 0.04 ($p < 8.2\times10^{-5}$) | 0.07 ± 0.05 ($p < 0.003$) | 0.07 ± 0.03 ($p < 0.008$) |
| Beta band ($Coherence^2$) | 0.08 ± 0.05 | 0.39 ± 0.1 ($p < 1.4\times10^{-30}$) | 0.21 ± 0.08 ($p < 5\times10^{-11}$) | 0.16 ± 0.08 ($p < 1.9\times10^{-6}$) | 0.16 ± 0.07 ($p < 3.4\times10^{-6}$) | 0.17 ± 0.06 ($p < 3.9\times10^{-7}$) |

*Mean ± 1 standard deviation. p-values as a comparison to "Rest".*

**Table S8 | Magnitude of $Coherence^2$ of bilateral delta band, theta band, alpha band, and beta band at 0.01 Hz**

| **$Coherence^2$ at 0.01 Hz** | **Alert** | **Asleep** | **All Data** |
|---|---|---|---|
| Delta band ($Coherence^2$) | 0.68 ± 0.16 | 0.34 ± 0.23 ($p < 2.8\times10^{-11}$) | 0.54 ± 0.22 ($p < 5.5\times10^{-5}$) |
| Theta band ($Coherence^2$) | 0.6 ± 0.19 | 0.59 ± 0.26 ($p < 0.67$) | 0.59 ± 0.25 ($p < 0.53$) |
| Alpha band ($Coherence^2$) | 0.62 ± 0.17 | 0.74 ± 0.13 ($p < 0.0007$) | 0.74 ± 0.13 ($p < 0.001$) |
| Beta band ($Coherence^2$) | 0.74 ± 0.12 | 0.8 ± 0.11 ($p < 0.08$) | 0.81 ± 0.1 ($p < 0.05$) |

*Mean ± 1 standard deviation. p-values as a comparison to "Alert".*

**Table S9 | Pearson's correlation coefficients of bilateral delta band, theta band, alpha band, and beta band**

| **Pearson's Correlation Coef.** | **Awake Rest** | **Whisking** | **Cont. NREM** | **Cont. REM** | **Alert** | **Asleep** | **All Data** |
|---|---|---|---|---|---|---|---|
| Delta band (R) | 0.15 ± 0.05 | 0.17 ± 0.08 ($p < 0.28$) | 0.17 ± 0.09 ($p < 0.27$) | 0.08 ± 0.05 ($p < 0.0002$) | 0.32 ± 0.08 ($p < 1.4\times10^{-14}$) | 0.19 ± 0.09 ($p < 0.03$) | 0.26 ± 0.1 ($p < 2.1\times10^{-8}$) |
| Theta band (R) | 0.14 ± 0.06 | 0.26 ± 0.09 ($p < 1.2\times10^{-8}$) | 0.15 ± 0.06 ($p < 0.68$) | 0.14 ± 0.07 ($p < 0.89$) | 0.3 ± 0.1 ($p < 5.7\times10^{-13}$) | 0.24 ± 0.1 ($p < 5.7\times10^{-7}$) | 0.28 ± 0.09 ($p < 8.4\times10^{-11}$) |
| Alpha band (R) | 0.09 ± 0.04 | 0.18 ± 0.08 | 0.2 ± 0.05 | 0.14 ± 0.05 | 0.25 ± 0.04 | 0.33 ± 0.04 | 0.31 ± 0.03 |

| | | | | | | | |
|---|---|---|---|---|---|---|---|
| | | ($p < 1.9\times10^{-6}$) | ($p < 6.2\times10^{-9}$) | ($p < 0.002$) | ($p < 8.7\times10^{-15}$) | ($p < 3\times10^{-24}$) | ($p < 1.5\times10^{-23}$) |
| Beta band (R) | 0.15 ± 0.06 | 0.14 ± 0.1 ($p < 0.45$) | 0.36 ± 0.06 ($p < 3.6\times10^{-23}$) | 0.25 ± 0.07 ($p < 5.5\times10^{-9}$) | 0.33 ± 0.08 ($p < 5.7\times10^{-19}$) | 0.46 ± 0.06 ($p < 4.3\times10^{-33}$) | 0.45 ± 0.06 ($p < 3.3\times10^{-33}$) |

*Mean ± 1 standard deviation. p-values as a comparison to "Rest".*

#### Table S10 | Coherence of Δ[HbT] vs. delta band, theta band, alpha band, and beta band at 0.1 Hz

| **Coherence$^2$ at 0.1 Hz** | **Awake Rest** | **Cont. NREM** | **Cont. REM** | **Alert** | **Asleep** | **All Data** |
|---|---|---|---|---|---|---|
| Δ[HbT]-Delta (Coherence$^2$) | 0.05 ± 0.04 | 0.07 ± 0.06 ($p < 0.15$) | 0.02 ± 0.02 ($p < 0.003$) | 0.06 ± 0.06 ($p < 0.52$) | 0.07 ± 0.06 ($p < 0.17$) | 0.06 ± 0.04 ($p < 0.21$) |
| Δ[HbT]-Theta (Coherence$^2$) | 0.01 ± 0.01 | 0.11 ± 0.06 ($p < 5.1\times10^{-20}$) | 0.06 ± 0.03 ($p < 6.2\times10^{-6}$) | 0.02 ± 0.02 ($p < 0.56$) | 0.06 ± 0.04 ($p < 1.3\times10^{-6}$) | 0.03 ± 0.03 ($p < 0.03$) |
| Δ[HbT]-Alpha (Coherence$^2$) | 0.02 ± 0.02 | 0.18 ± 0.05 ($p < 1.5\times10^{-37}$) | 0.04 ± 0.03 ($p < 0.009$) | 0.02 ± 0.02 ($p < 0.8$) | 0.07 ± 0.05 ($p < 2.1\times10^{-6}$) | 0.04 ± 0.02 ($p < 0.03$) |
| Δ[HbT]-Beta (Coherence$^2$) | 0.04 ± 0.03 | 0.28 ± 0.06 ($p < 4.3\times10^{-57}$) | 0.05 ± 0.04 ($p < 0.37$) | 0.04 ± 0.04 ($p < 0.82$) | 0.11 ± 0.04 ($p < 8.9\times10^{-11}$) | 0.09 ± 0.03 ($p < 1.2\times10^{-7}$) |
| Δ[HbT]-Gamma (Coherence$^2$) | 0.18 ± 0.11 | 0.32 ± 0.09 ($p < 7.9\times10^{-10}$) | 0.08 ± 0.07 ($p < 1.8\times10^{-5}$) | 0.27 ± 0.14 ($p < 6\times10^{-5}$) | 0.28 ± 0.14 ($p < 7.6\times10^{-6}$) | 0.28 ± 0.13 ($p < 4.1\times10^{-6}$) |

*Mean ± 1 standard deviation. p-values as a comparison to "Rest".*

#### Table S11 | Coherence of Δ[HbT] vs. delta band, theta band, alpha band, and beta band at 0.01 Hz

| **Coherence at 0.01 Hz** | **Alert** | **Asleep** | **All Data** |
|---|---|---|---|
| Δ[HbT]-Delta (Coherence) | 0.1 ± 0.11 | 0.14 ± 0.12 ($p < 0.05$) | 0.2 ± 0.11 ($p < 2\times10^{-5}$) |
| Δ[HbT]-Theta (Coherence) | 0.11 ± 0.1 | 0.5 ± 0.24 ($p < 7.9\times10^{-14}$) | 0.42 ± 0.19 ($p < 1.2\times10^{-10}$) |
| Δ[HbT]-Alpha (Coherence) | 0.14 ± 0.13 | 0.39 ± 0.19 ($p < 2\times10^{-11}$) | 0.43 ± 0.14 ($p < 4.2\times10^{-15}$) |
| Δ[HbT]-Beta (Coherence) | 0.16 ± 0.18 | 0.52 ± 0.18 ($p < 1.7\times10^{-16}$) | 0.55 ± 0.13 ($p < 7.9\times10^{-19}$) |
| Δ[HbT]-Gamma (Coherence) | 0.38 ± 0.18 | 0.46 ± 0.16 ($p < 0.06$) | 0.51 ± 0.14 ($p < 0.001$) |

*Mean ± 1 standard deviation. p-values as a comparison to "Alert".*

#### Table S12 | Model OOB error vs. shuffled-date model OOB error for each IOS animal

| **Animal ID** | **Random Forest Out-of-bag (OOB) error (%)** | **Mean OOB (%) of 100 shuffled data models** |
|---|---|---|
| ***T99*** | 7.5 | 47.1 |
| ***T101*** | 7.2 | 32.8 |
| ***T102*** | 6.3 | 33.0 |
| ***T103*** | 5.6 | 25.8 |
| ***T105*** | 6.0 | 31.3 |
| ***T108*** | 8.1 | 45.3 |
| ***T109*** | 4.6 | 22.6 |
| ***T110*** | 5.2 | 20.7 |
| ***T111*** | 8.3 | 39.8 |

| | | |
|---|---|---|
| ***T119*** | 7.5 | 39.8 |
| ***T120*** | 8.7 | 52.1 |
| ***T121*** | 9.3 | 53.4 |
| ***T122*** | 7.5 | 35.1 |
| ***T123*** | 7.2 | 26.2 |

#### *Fig. S1 - Turner et al. 2020*

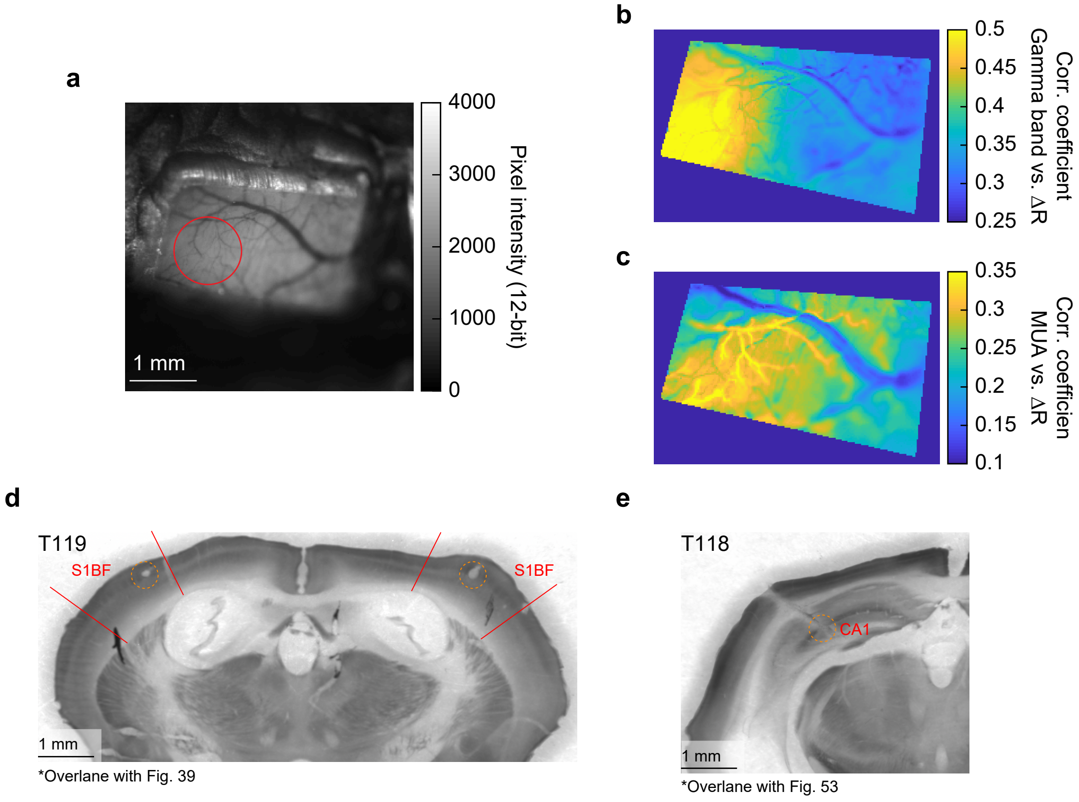


*The Mouse Brain in Stereotaxic Coordinates, 3rd Edition (Franklin & Paxinos)

*Fig. S2 - Turner et al. 2020*

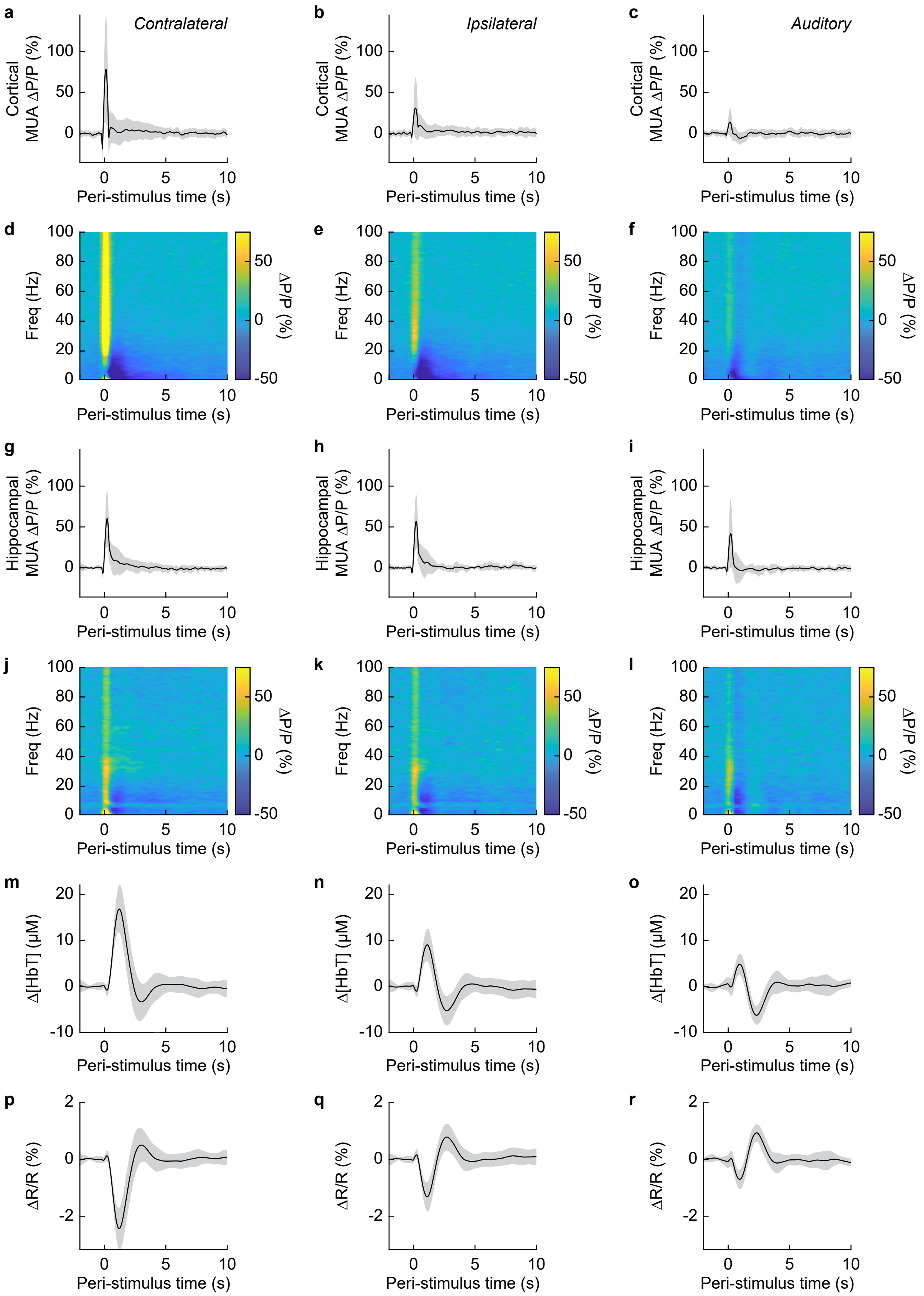

Fig. S3 - Turner et al. 2020

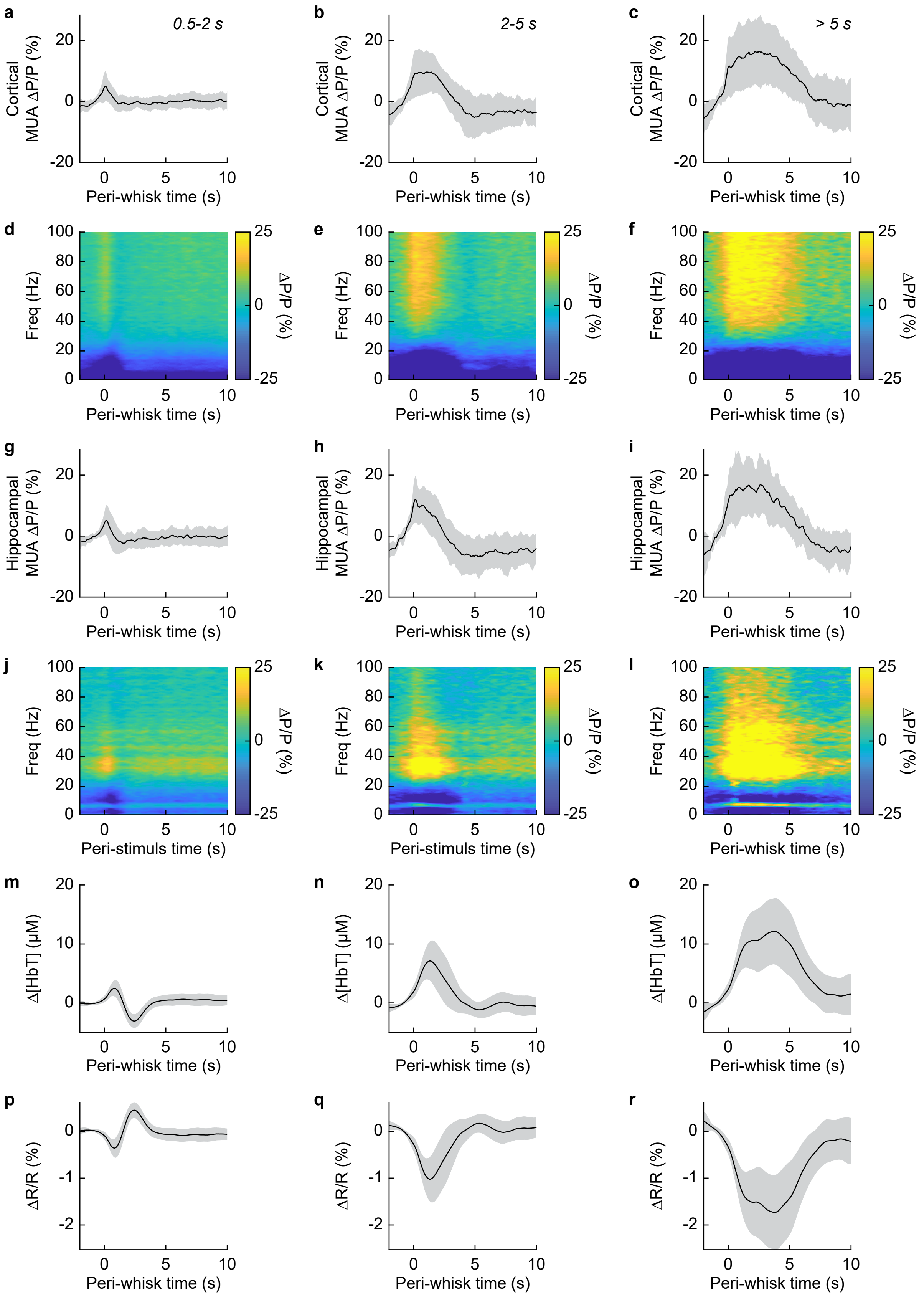

***Fig. S4 - Turner et al. 2020***

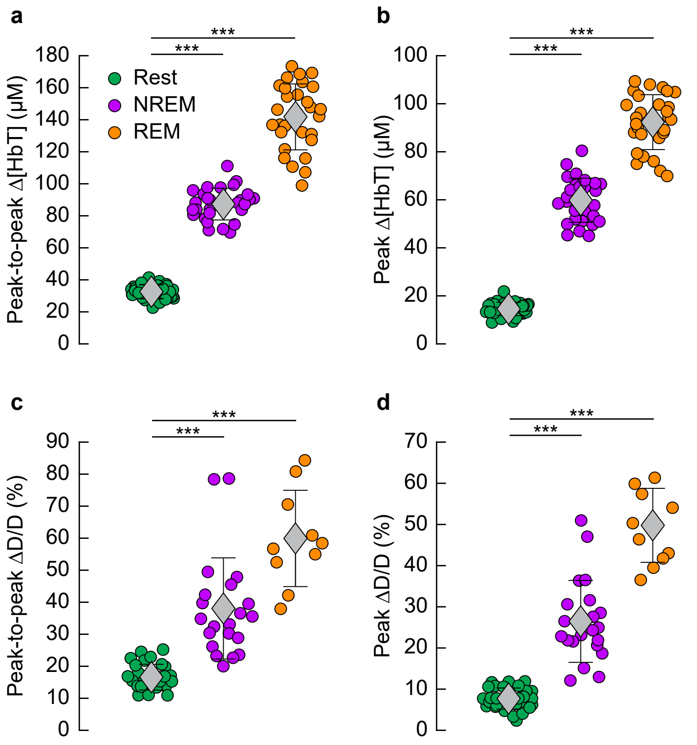

***Fig. S5 - Turner et al. 2020***

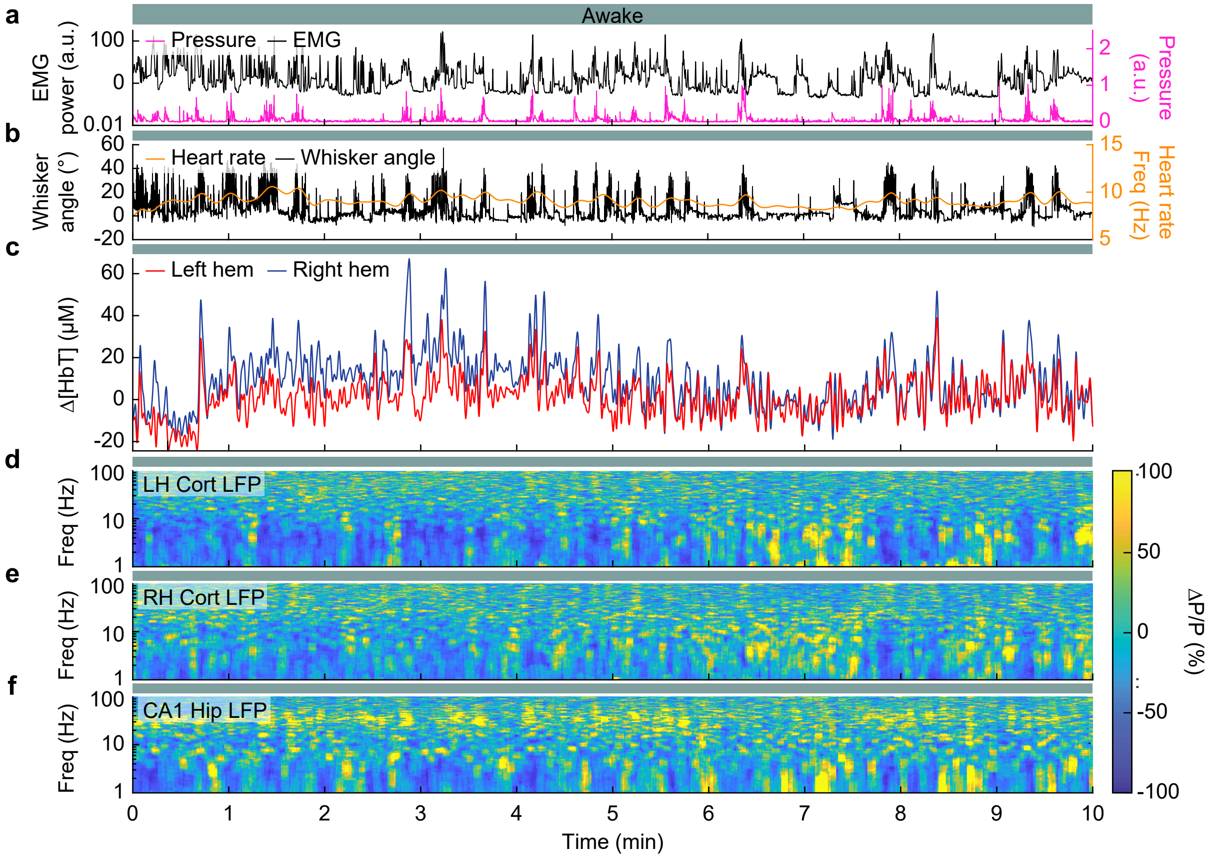

*Fig. S6 - Turner et al. 2020*

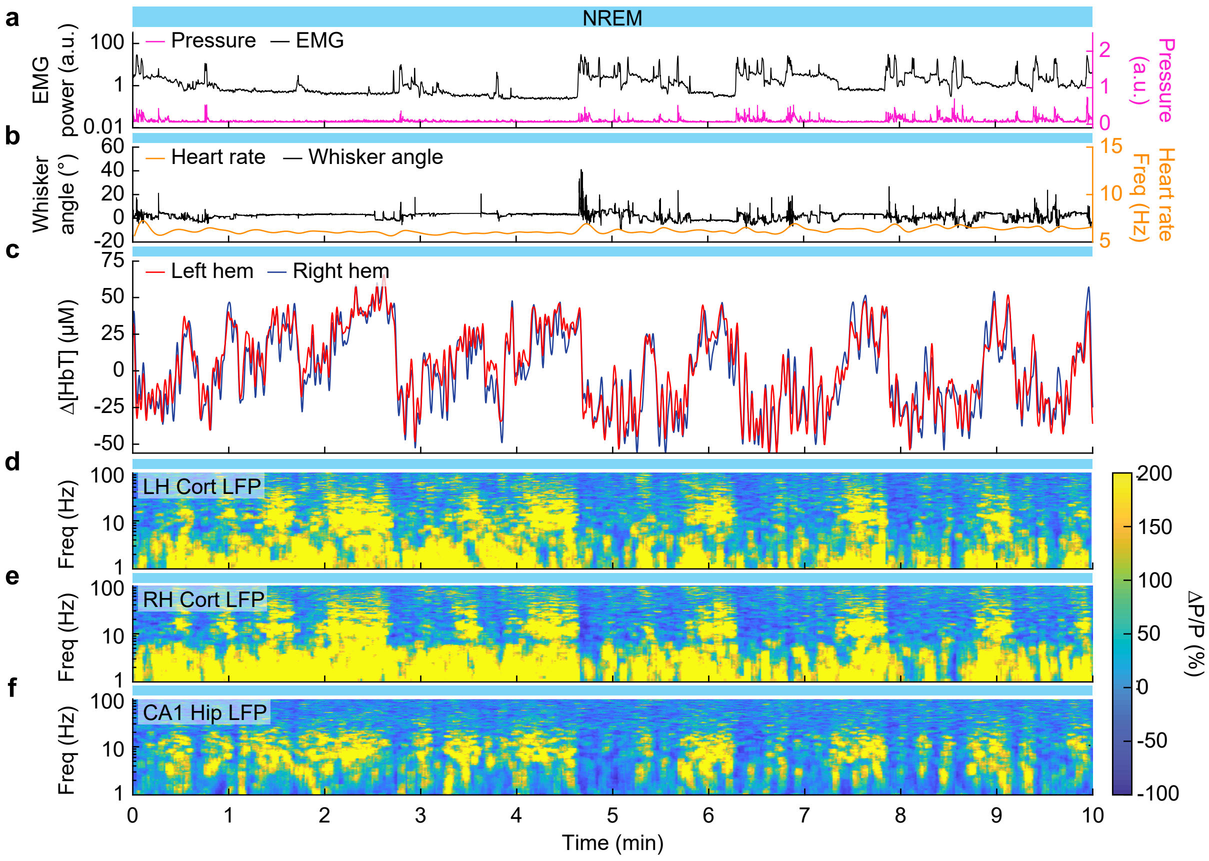

*Fig. S7 - Turner et al. 2020*

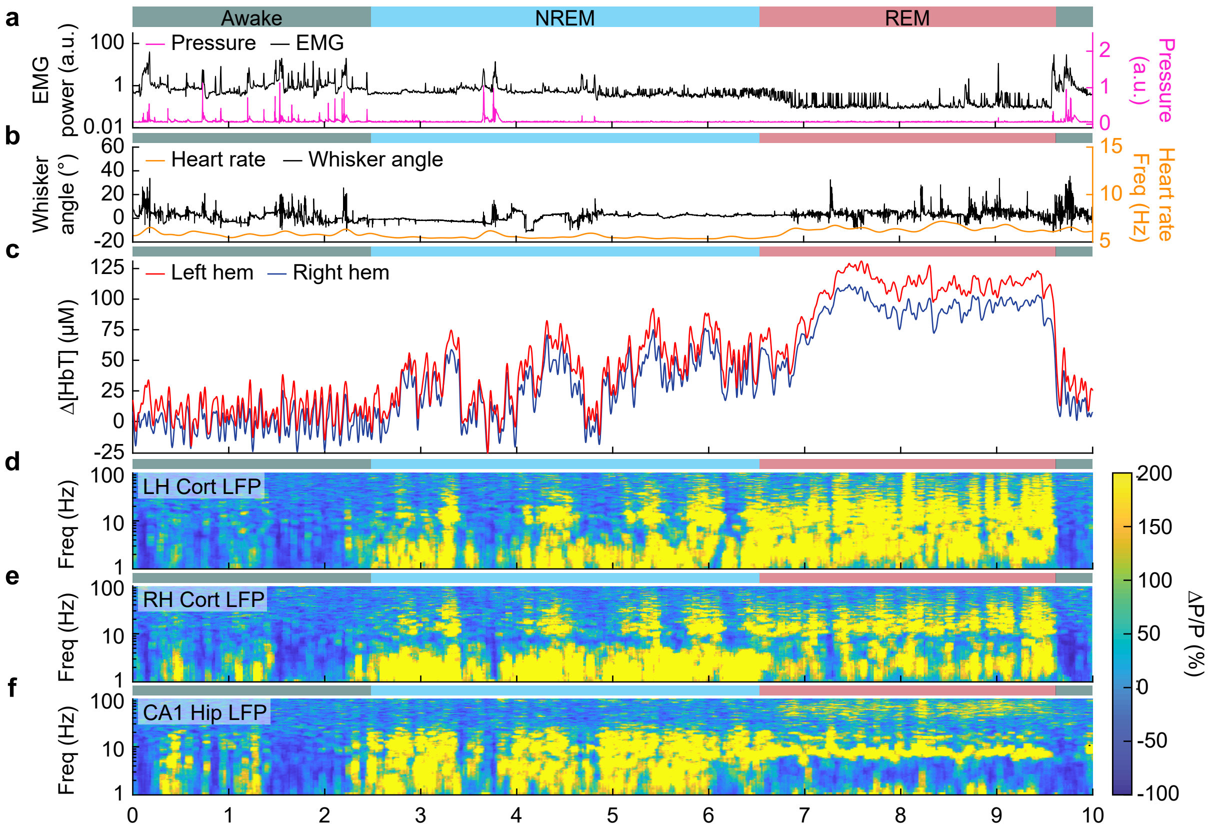

*Fig. S8 - Turner et al. 2020*

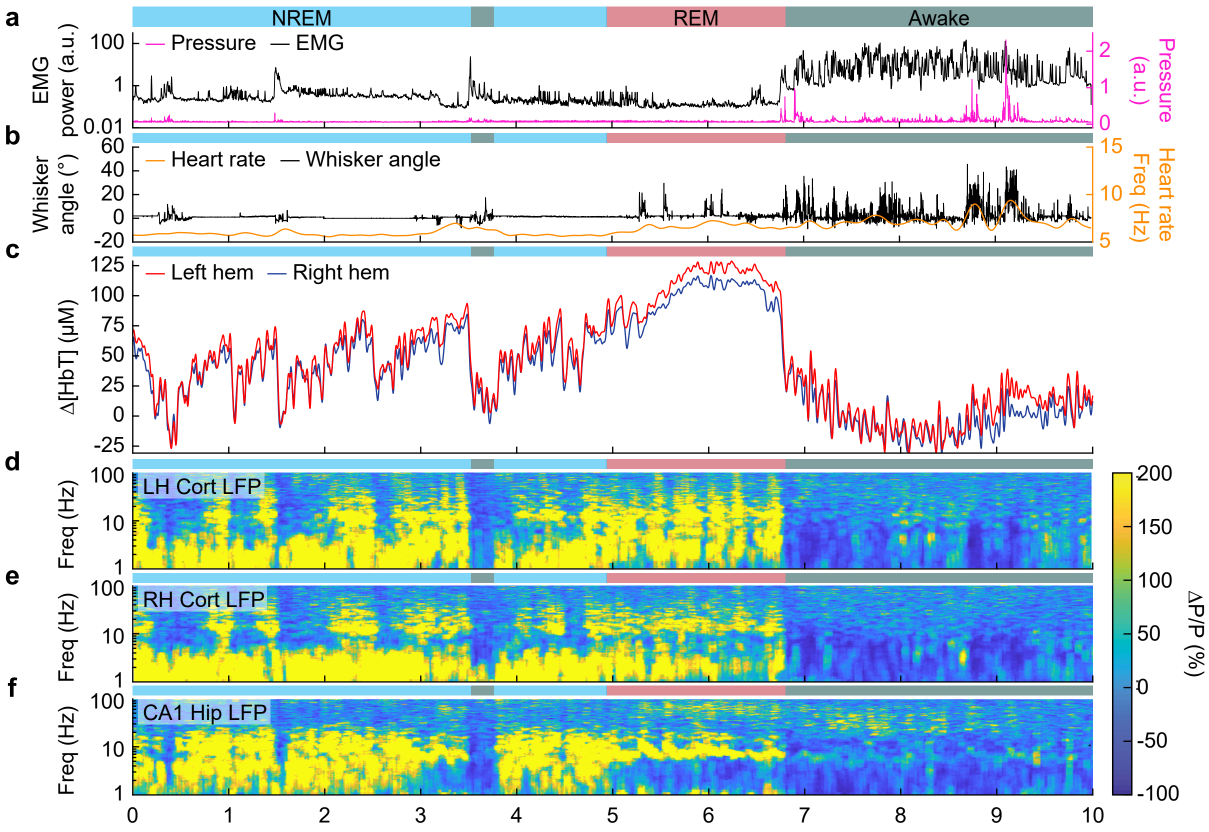

*Fig. S9 - Turner et al. 2020*

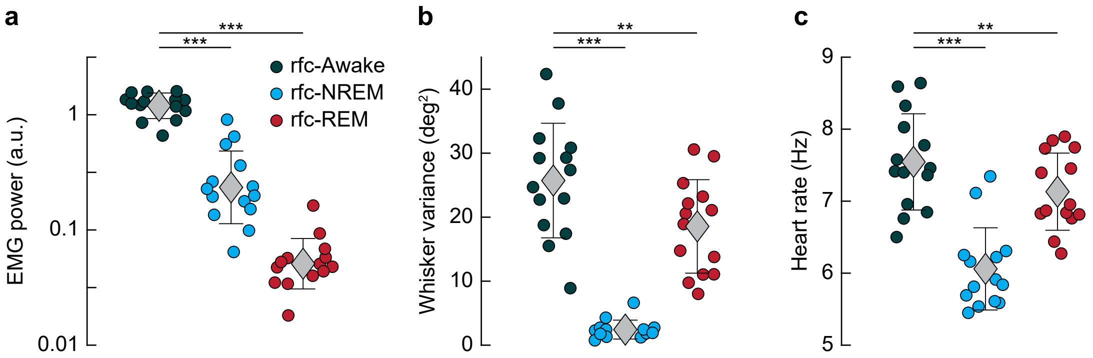

*Fig. S10 - Turner et al. 2020*

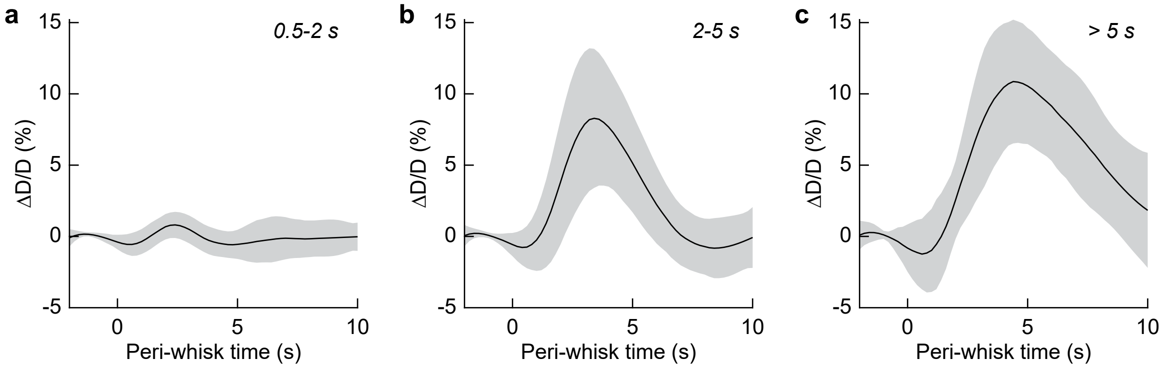

*Fig. S11 - Turner et al. 2020*

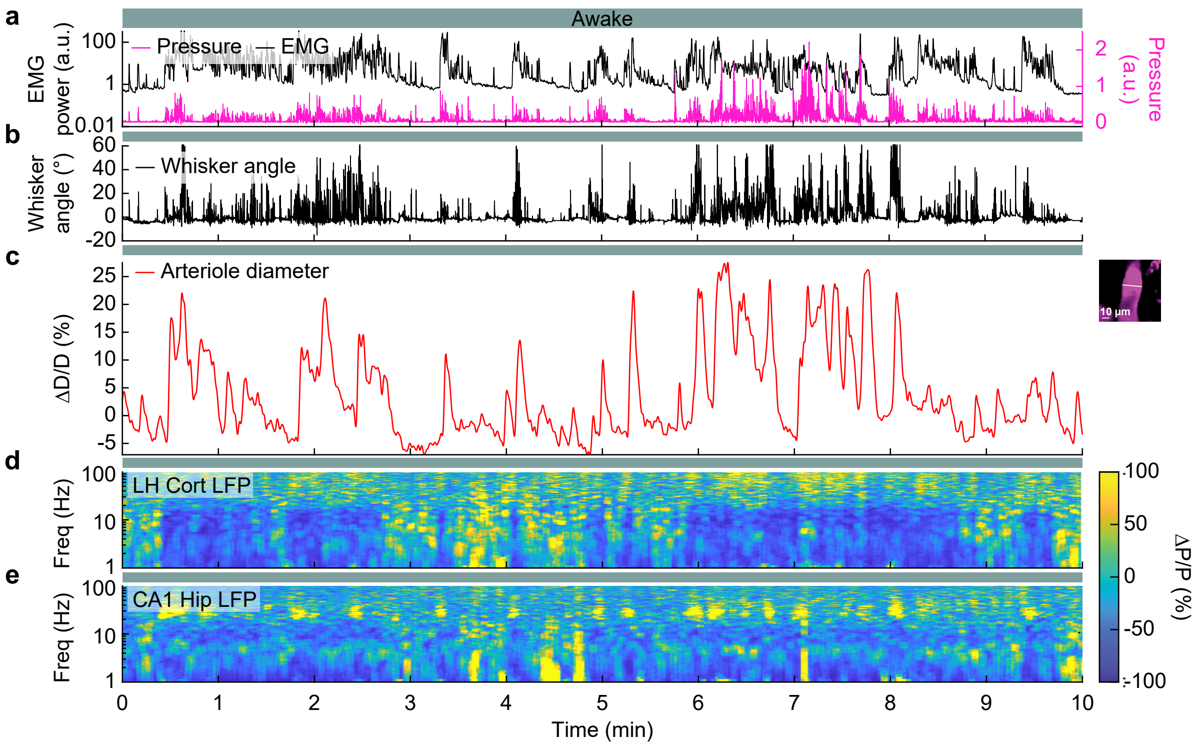

*Fig. S12 - Turner et al. 2020*

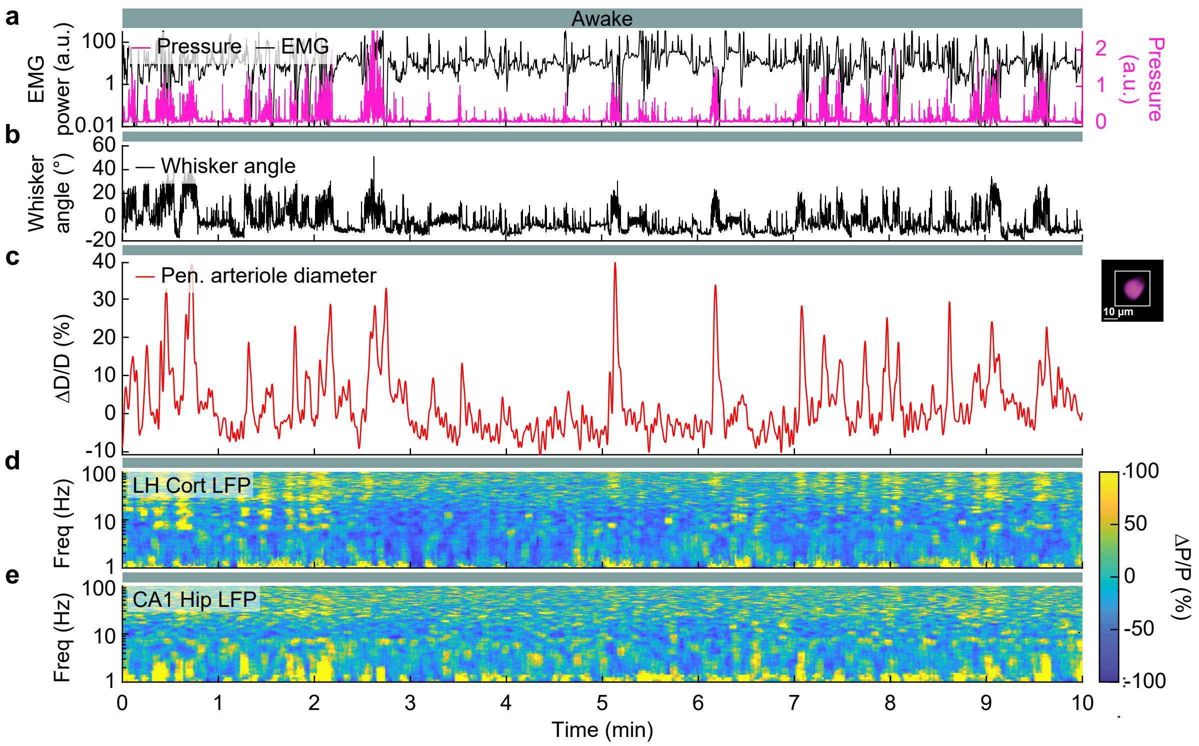

*Fig. S13 - Turner et al. 2020*

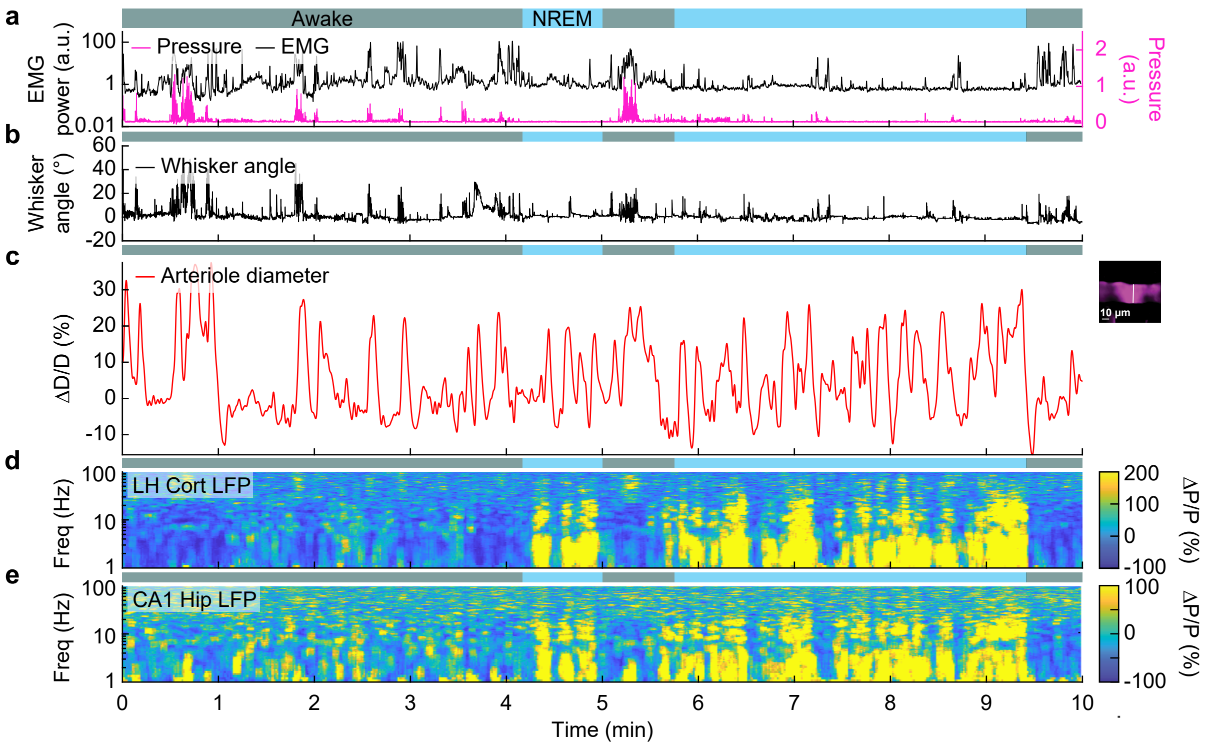

*Fig. S14 - Turner et al. 2020*

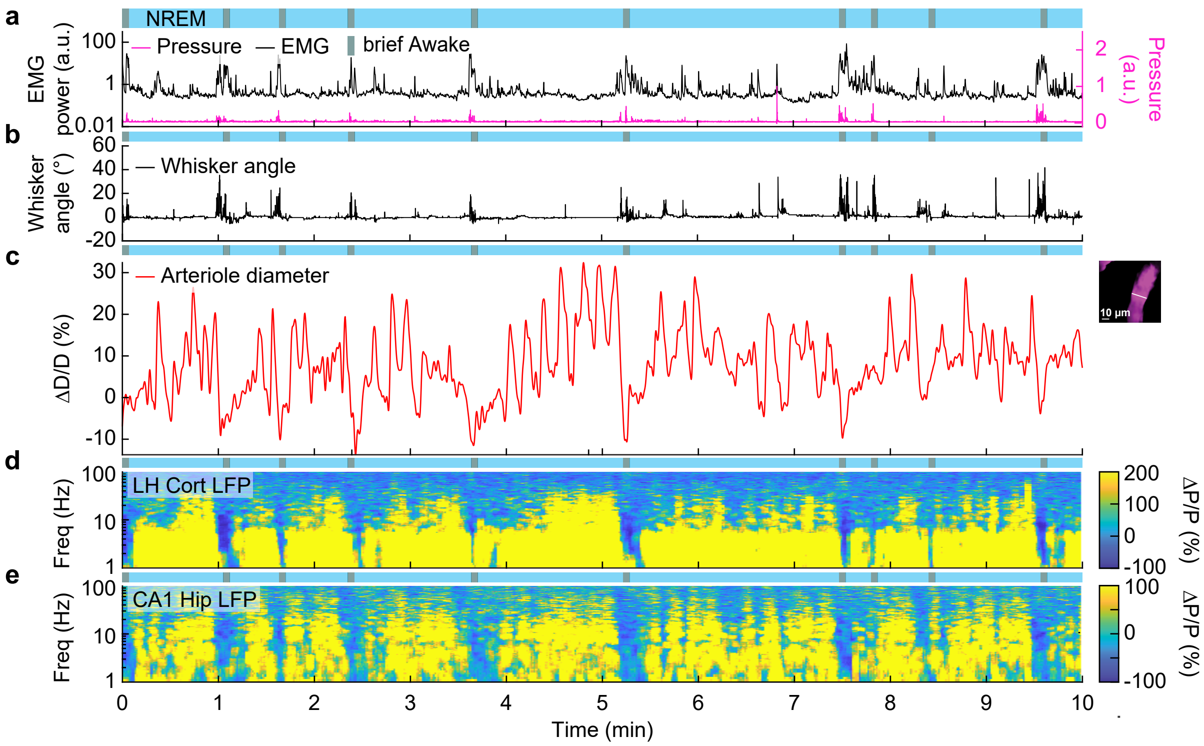

***Fig. S15 - Turner et al. 2020***

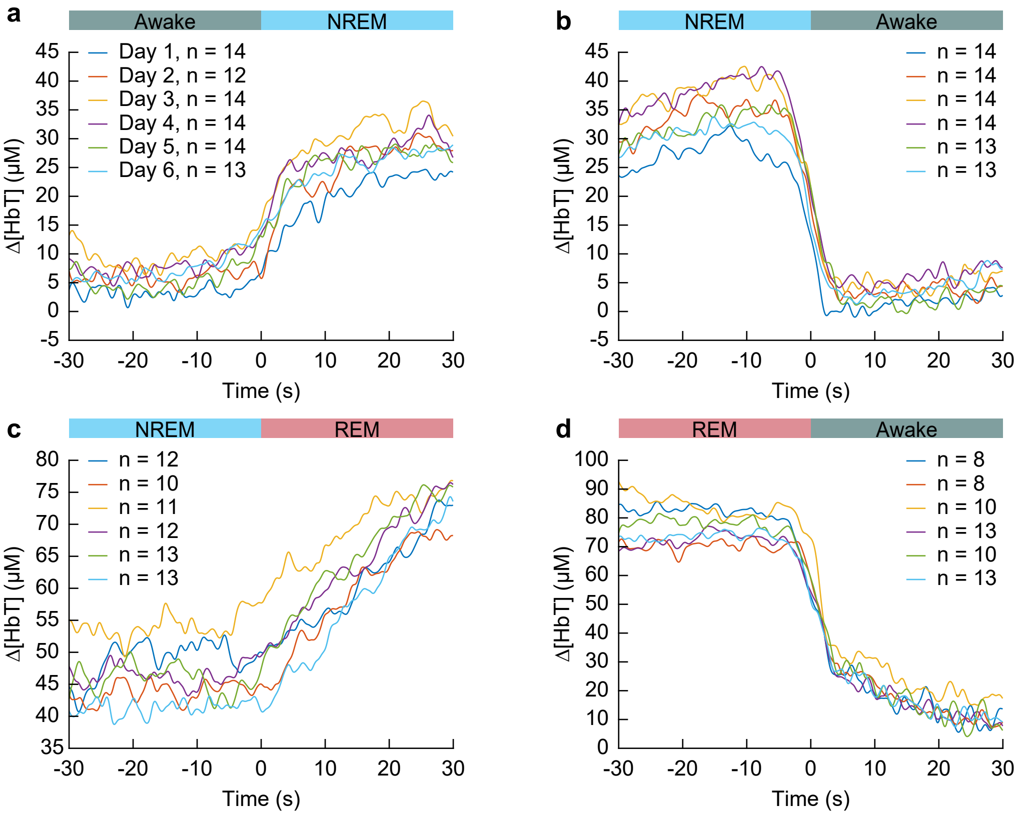

*Fig. S16 - Turner et al. 2020*

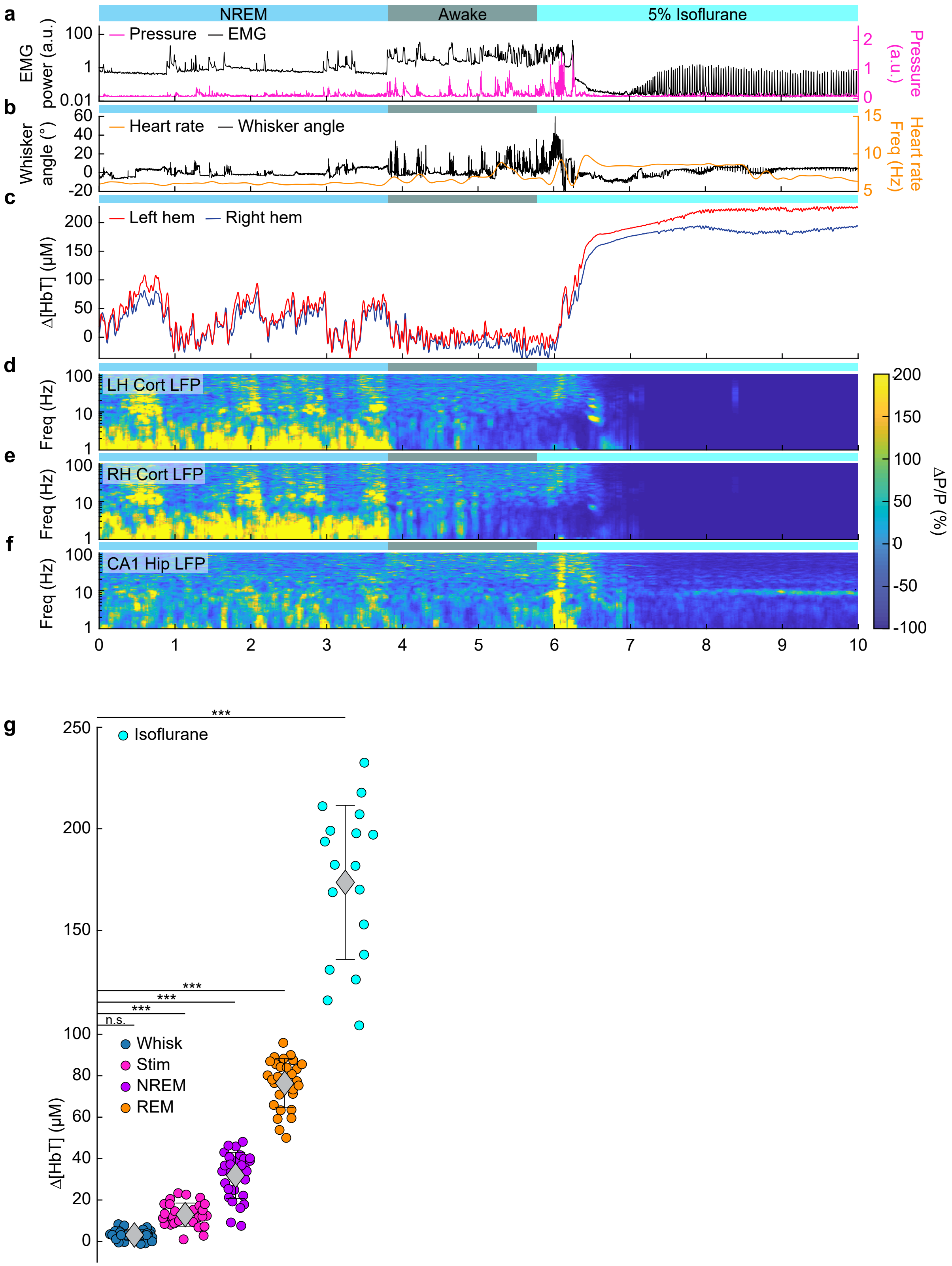

### *Fig. S17 - Turner et al. 2020*

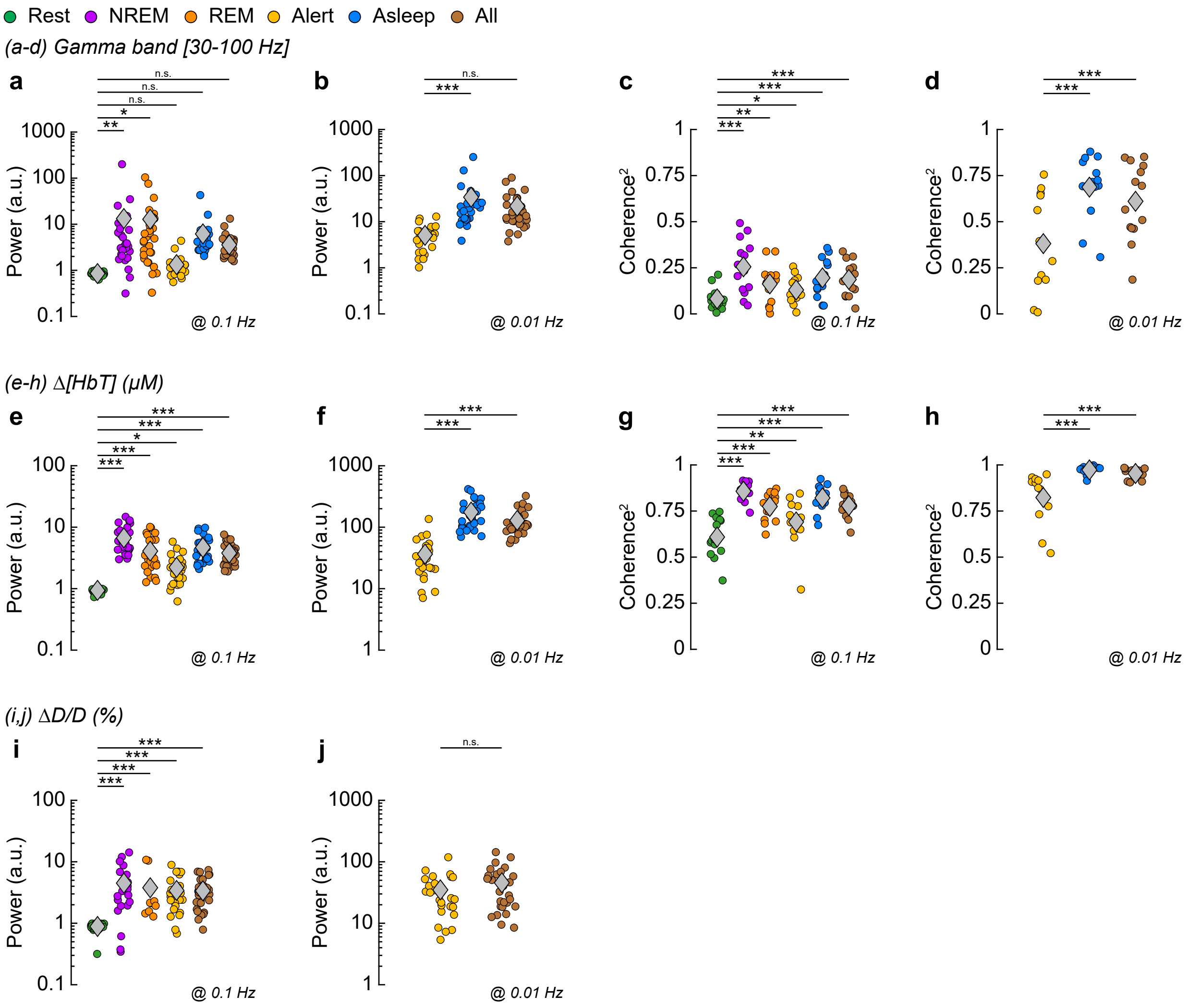

***Fig. S18 - Turner et al. 2020***

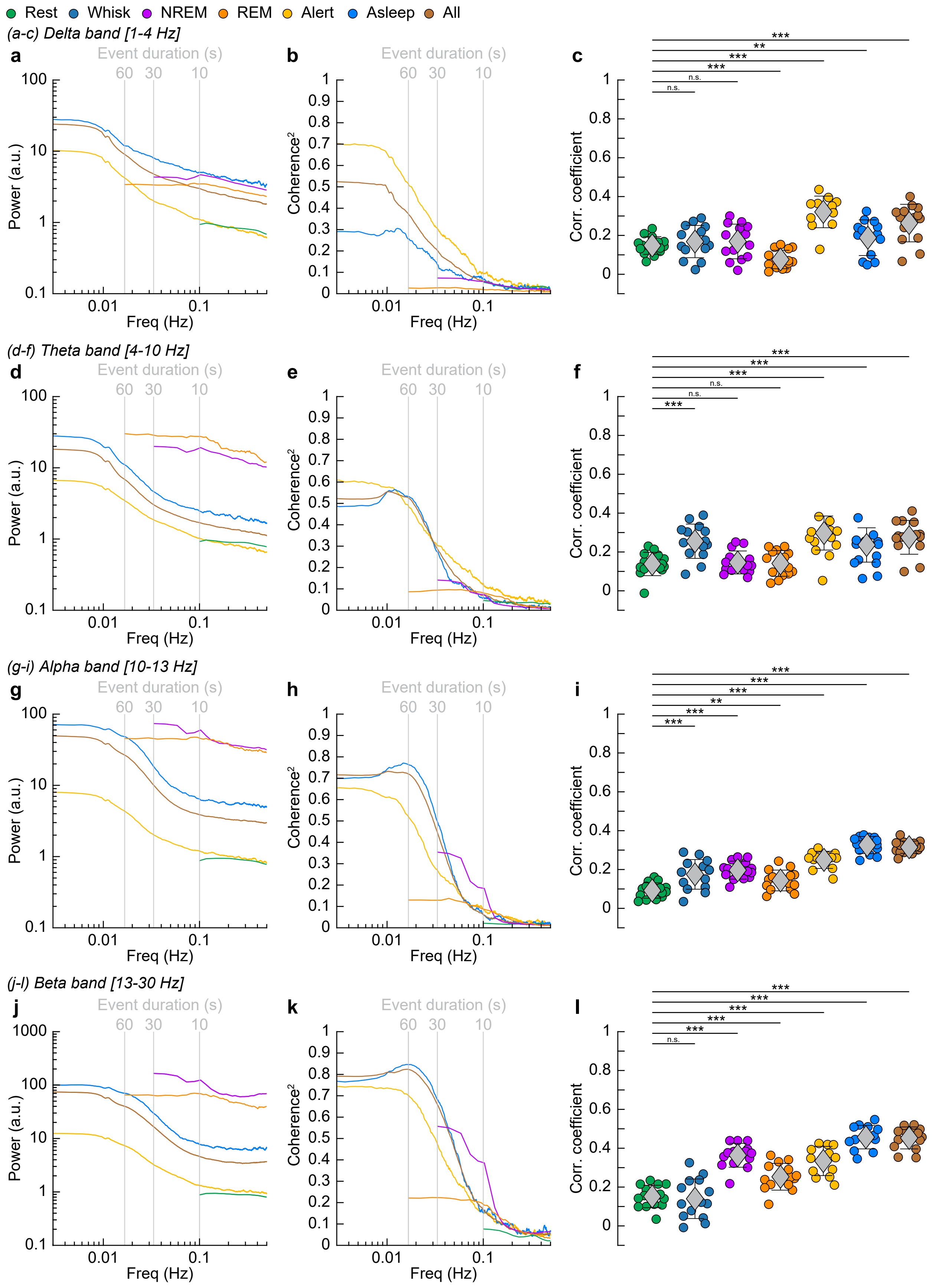

**Fig. S19 -** *Turner et al. 2020*

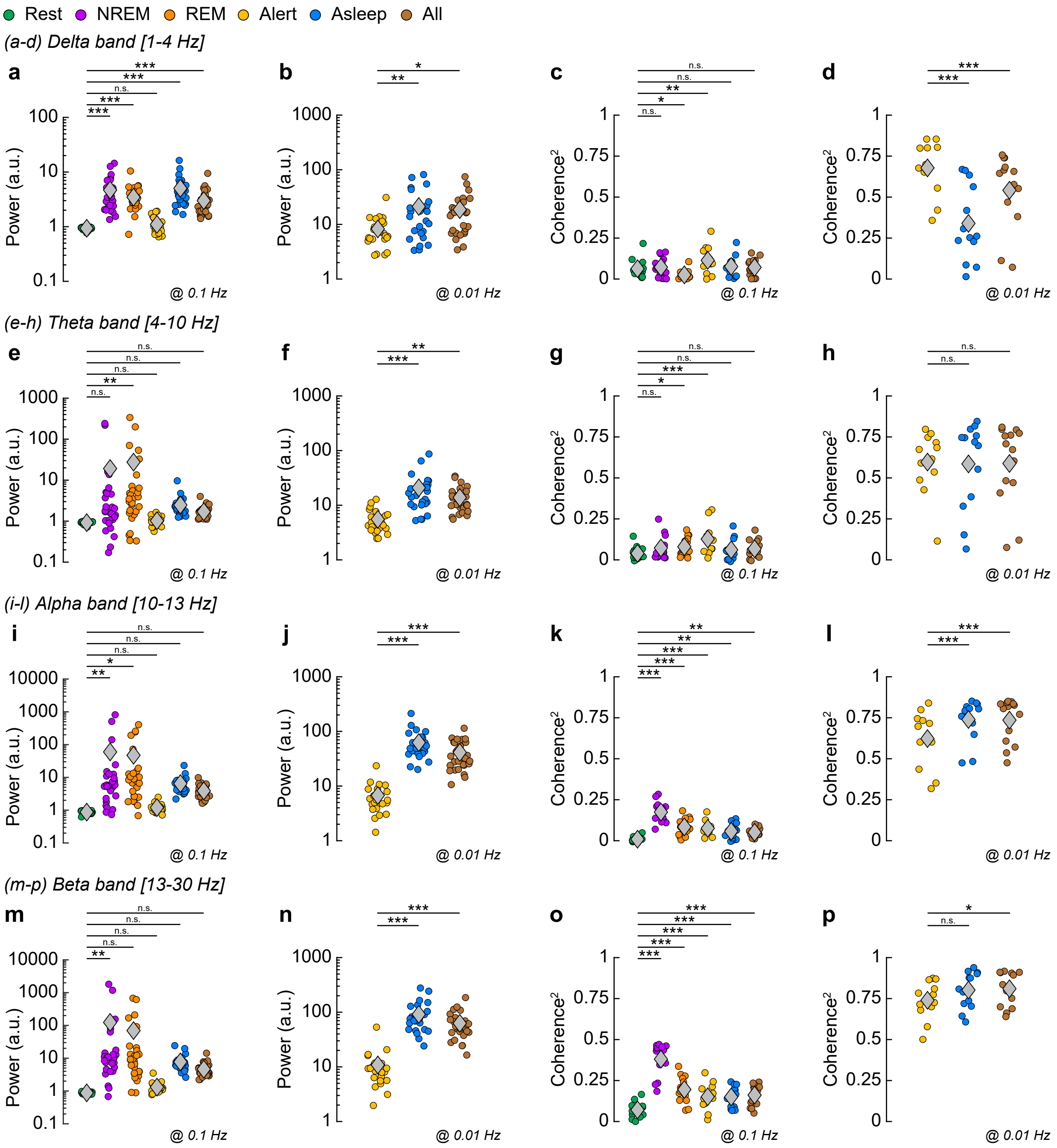

#### *Fig. S20 - Turner et al. 2020*

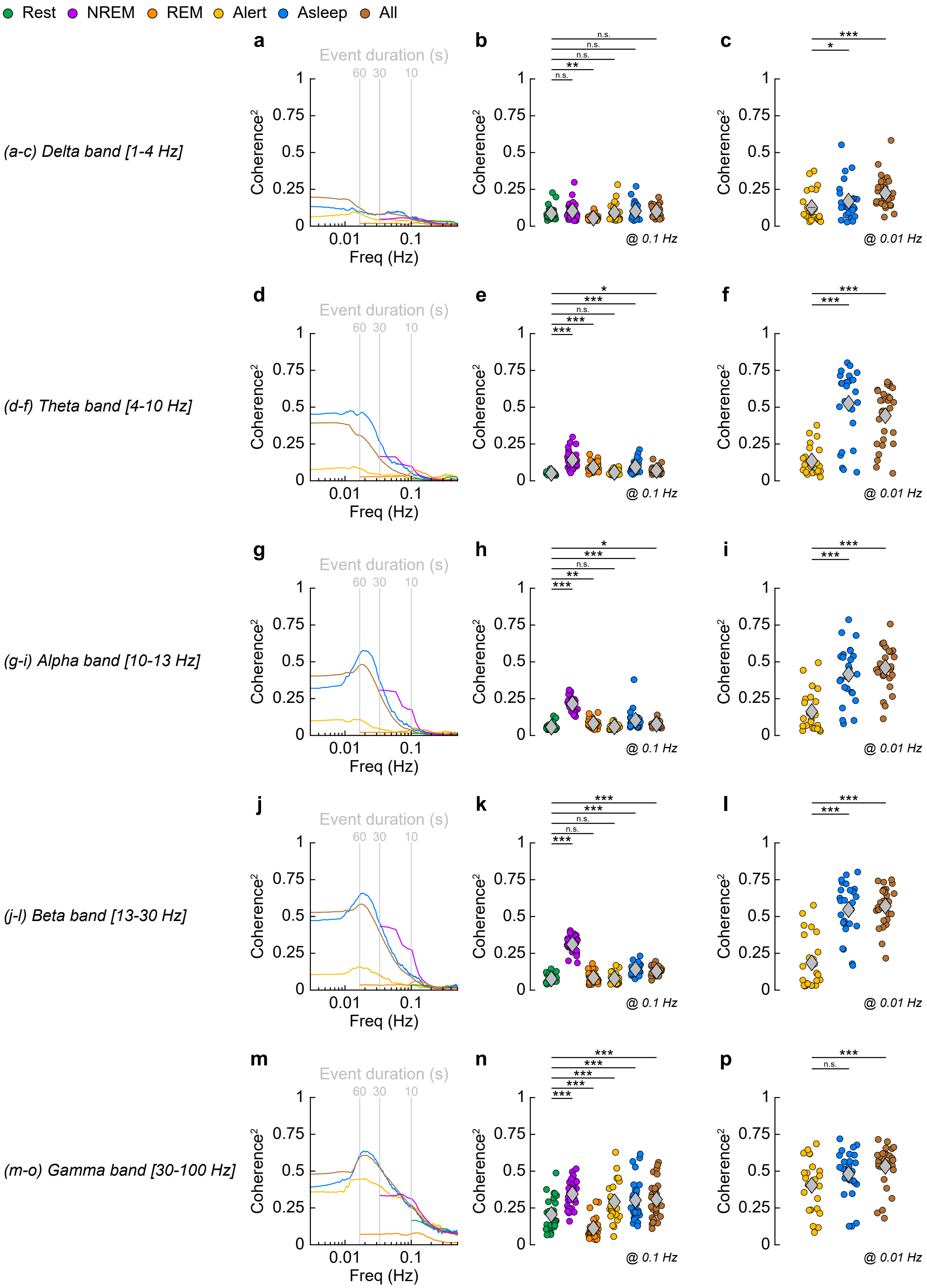

*Fig. S21 - Turner et al. 2020*

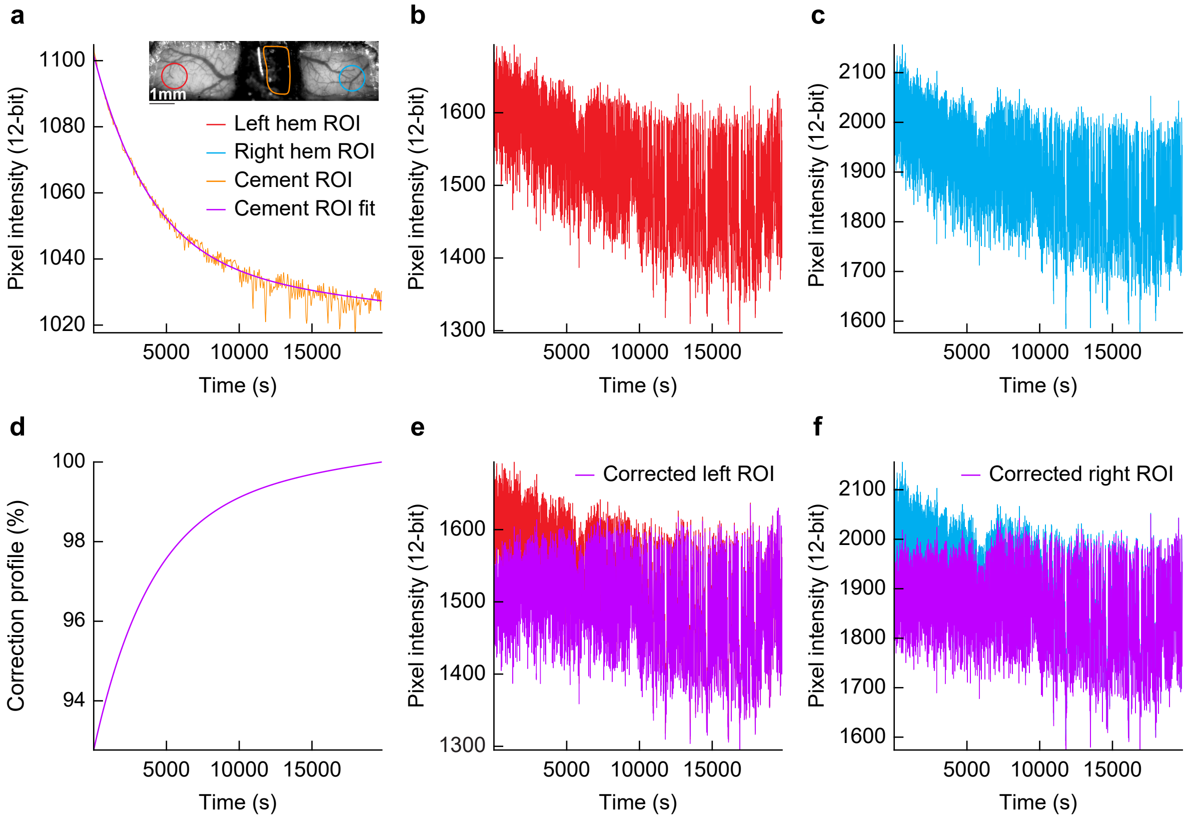

**Fig. S22 -** *Turner et al. 2020*

**a**

| True Class \ Predicted Class | rfc-Awake | rfc-NREM | rfc-REM | | |
|---|---|---|---|---|---|
| **rfc-Awake** | 19066 | 1383 | 15 | 93.2% | 6.8% |
| **rfc-NREM** | 1347 | 14350 | 319 | 89.6% | 10.4% |
| **rfc-REM** | 55 | 234 | 1751 | 85.8% | 14.2% |
| | 93.2% | 89.9% | 84.0% | | |
| | 6.8% | 10.1% | 16.0% | | |

**Total accuracy: 91.3 (%)**
